# Wheel-running exercise during adolescence may not convincingly impact cocaine conditioned place preference in male C57BL/6J mice

**DOI:** 10.1101/477299

**Authors:** Louis-Ferdinand Lespine, Ezio Tirelli

## Abstract

Wheel-running in rodents can mitigate addiction-related effects of drugs of abuse like cocaine. However, experiments using conditioned place preference (CPP) are conflicting, warranting further studies. Our purpose was to test whether wheel-running during adolescence could impact the formation and long-term retention of CPP to cocaine in mice. Male C57BL/6J mice were individually housed either with (n=32) or without (n=32) a running wheel from the age of 35 days. Behavioral testing began 3 weeks after such housing, mice underwent a baseline session followed by 10 once-daily conditioning sessions receiving peritoneal injections of 10 mg/kg cocaine and saline on alternate days (n=16), control mice receiving saline every day (n=16). One and 21 days after the last conditioning session, they were tested for CPP. Both groups exhibited comparable well-marked cocaine-induced CPP in both post-conditioning tests resulting in a negligible interaction between housing and the pharmacological treatment (η²p < 0.01). These results, along with the discrepancy found in the literature, question the nature (and the robustness) of the effects that exercise induces on CPP to cocaine.

---

Animal research has shown that rodents housed with a running wheel (a model of aerobic exercise) exhibited a reduced motivation for self-administered various addictive drugs including cocaine, as well as attenuated initiation or expression of psychomotor sensitization to that drug [1-3]. However, studies using conditioned place preference (CPP) provide conflicting results. Although rats housed with a wheel displayed lower CPP to morphine compared to their sedentary counterparts [4], accentuated CPP to morphine and cocaine have been reported in exercising rats and mice [5-7]. Intermediately, other studies have observed a negligible difference in CPP levels induced by cocaine or morphine between exercising and sedentary rodents [8,9]. In addition to procedural differences between experiments, aspects of the methods used in some studies such as the absence of a baseline pre-conditioning session or a control group in the design or, when present, in the data analyses, make the whole picture of results difficult to understand. In the present study, we intended to test whether wheel-running was able to impact the formation of CPP to cocaine in male C57BL/6J mice by using a factorial 2 × 2 design with mice housed either with or without a wheel and receiving either cocaine or saline injections to establish CPP.

Male C57BL/6J mice, obtained at 28 days of age from JANVIER, Le-Genest-Saint-Isle, France, were housed upon arrival in groups of eight for one week in large transparent polycarbonate cages (38.2 × 22 cm surface × 15 cm height). After that acclimation period and until the end of experimentation, these 35-day-old mice were randomly assigned to exercise (n=32) or sedentary conditions (no wheel, n=32), each group receiving cocaine or saline during testing (n=16), the assignment into groups being based on a computer-generated randomization schedule. The determination of the sample size was informal and based on the median number of animals used in the previous experiments mentioned above (n=8 per group) that we elected to double. Thus, we had 80% power at an alpha level of 5% to detect a partial eta-squared (η²p) of 0.11 (intermediate-to-large effect) reflecting the importance of the interaction between housing and the pharmacological treatment on CPP.

A running wheel consists of an orange polycarbonate saucer-shaped disk (diameter 15 cm, circumference 37.8 cm), which allowed an open running surface, mounted on a plastic cup-shaped base (height 4.5 cm) via a bearing pin so as to being inclined from the vertical plane at an angle of 35° (ENV-044, Med Associates; St Albans, VT, USA). The base was fixed on a transparent acryl-glass plate. Running was monitored and recorded continuously using a wireless system, each wheel being connected to a USB interface hub (DIG-804, Med Associates) which relayed data to a Wheel Manager Software (SOF-860, Med Associates). Mice were individually housed in TECHNIPLAST transparent polycarbonate cages (32.5 } 17 cm surface × 14 cm height) with pine sawdust bedding, tap water, and food (standard pellets, CARFIL QUALITY, Oud-Turnhout, Belgium) being continuously available. The animal room was maintained on a 12:12 h dark-light cycle (lights on at 7:00 am), at an ambient temperature of 20-24°C.

(−)-Cocaine hydrochloride (BELGOPIA, Louvain-La-Neuve, Belgium), dissolved in an isotonic saline solution (0.9% NaCl), was injected at a dose of 10 mg/kg in a volume of 0.01 ml/g of body weight, based on previous studies in which the effects of free exercise on cocaine-induced CPP were investigated [6,7,9]. The control treatment consisted of an equal volume of isotonic saline solution. All injections were made via the peritoneal route as was the case in the studies mentioned above.

The testing apparatus consisted of a battery of eight place preference stations (Opto-Max Activity Meter v2.16; Columbus Instruments, Columbus, OH), each station comprising two equally-sized compartments (21 × 20 cm surface × 20 cm height) separated by a removable guillotine door. One compartment comprised four black walls with a smooth black floor and the other four white walls with an embossed white floor (these floors consisted of removable clear acrylic-glass panels). The time spent (in seconds) within each compartment were automatically recorded and analyzed by an interface (Opto-Max) via an array of two horizontal sensors mounted alongside opposing lengths (seven infrared beams at one inch-intervals corresponding to 2.54 cm).

Behavioral testing involved the following four phases (see Fig. 1). (1) During the baseline pre-conditioning session (1^st^ session), all mice were injected with saline and placed in the intermediate zone of the two compartments with free access to them during 20 min, the time spent within each compartment being recorded. (2) The conditioning phase lasted 10 consecutive once-daily sessions and involved alternate peritoneal injections of cocaine or saline (2^nd^ to 11^th^ sessions). Mice were injected immediately before being placed into the test chamber, the monitoring beginning as early as an infrared beam was interrupted. Due to the use of a biased apparatus (natural avoidance of the white compartment), we used a non-counterbalanced assignment procedure in which the non-preferred compartment was associated with the drug effects [10]. On the five odd conditioning days (1, 3, 5, 7 and 9), mice were injected with cocaine and confined within the drug-paired compartment (initially non-preferred) for 20 min. On the five even conditioning days (2, 4, 6, 8 and 10), mice were injected with saline and confined for 20 min in the opposite compartment. Control animals received saline in all sessions. (3) The third phase consisted of a CPP test which took place 24 h after the last conditioning day (12^th^ session) on which all groups were injected with saline and placed in the intermediate zone of the two compartments with free access to both black and white compartments for 20 min (as for the pre-conditioning session). (4) The last phase consisted of a long-term retention CPP test which took place 21 days after the last conditioning day (13^th^ session), the procedure remaining the same.

**Fig. 1.**
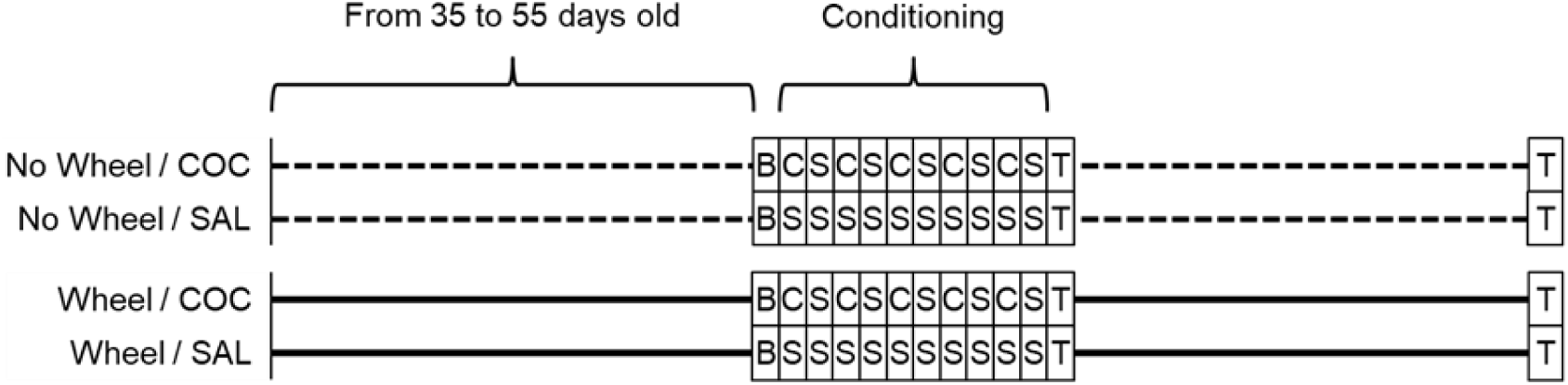
Experimental timeline and experimental design. At 35 days of age, mice were individually housed either in the presence or the absence of a running wheel (“Wheel” and “No Wheel” groups, n=32) and were left in these conditions until the end of experimentation. Mice from the two types of housing condition were injected peritoneally with either saline (SAL) or 10 mg/kg cocaine (COC) forming the four experimental groups of the present study with n=16. B: baseline pre-conditioning session under saline where animals had free access to the entire apparatus; C: cocaine administration associated with the initially non-preferred compartment; S: saline injection associated with the initially preferred compartment. T: CPP test under saline where animals had free access to the entire apparatus performed 24 h and 21 days after the last conditioning session.

All mice were familiarized with handling and injected twice with a saline solution in the animal room during the preceding week of testing. All experimental conditions were systematically represented by two mice within each of the eight once-daily 20-min sessions. Therefore, a 20-min session formed a block including eight mice individually tested in as many test chambers (randomized block design). After each 20-min session, animals were returned to the animal room within 10 min, and the test chambers were cleaned with a disinfectant. All procedures were conducted between 9:00 am and 12:30 pm.

All experimental treatments and animal maintenance were reviewed by the University of Liège Animal Care and Experimentation Committee (animal subjects review board), which gave its approval according to the Belgian implementation of the animal welfare guidelines laid down by the European Union (“Arrêté Royal relatif à la protection des animaux d’expérience” released on 23 May 2013, and “Directive 2010/63/EU of the European Parliament and of the Council of 22 September 2010 on the protection of animals used for scientific purposes”).

The CPP score consisted of the absolute difference between the pre- and post-conditioning sessions (by subtracting the amount of time spent in the drug-paired side before conditioning from the amount of time spent in the drug-paired side after conditioning). These scores were submitted to a fixed-model 2 × 2 ANOVA with the housing condition and the pharmacological treatment as between-group factors and time of the day as a blocking factor (8 levels). The basic effect of cocaine was confirmed for each housing condition by planned comparisons as one-tailed *t*-tests. The effect size of the ANOVAs main effects and interactions, and planned comparisons were estimated and given by ƞ²p and Cohen’s *d* respectively, conventionally classifiable as small (0.01 or 0.20), medium (0.06 or 0.50) or large (0.14 or 0.80). Assumptions of normality, homogeneity of variances and sphericity (in the case of repeated measures) were verified via Shapiro-Wilk, Levene and Mauchly’s tests, respectively. In the case of significant Levene’s test, raw data were subjected to a square-root transformation to more nearly meet the assumption of homogeneity of variances. For the sake of clarity, raw values (means ± SEM) are presented in the figures. The statistical significance threshold was set at the conventional alpha-level of 0.05.

As presented in Fig. 2, mice showed a rapid increase in wheel-running over the two first weeks until reaching a mean of 16.11 km (± 0.69 km SEM) on the 18^th^ day (peak of activity). During the following weeks, wheel-running plateaued and remained constant until the end of the experiment.

**Fig. 2.**
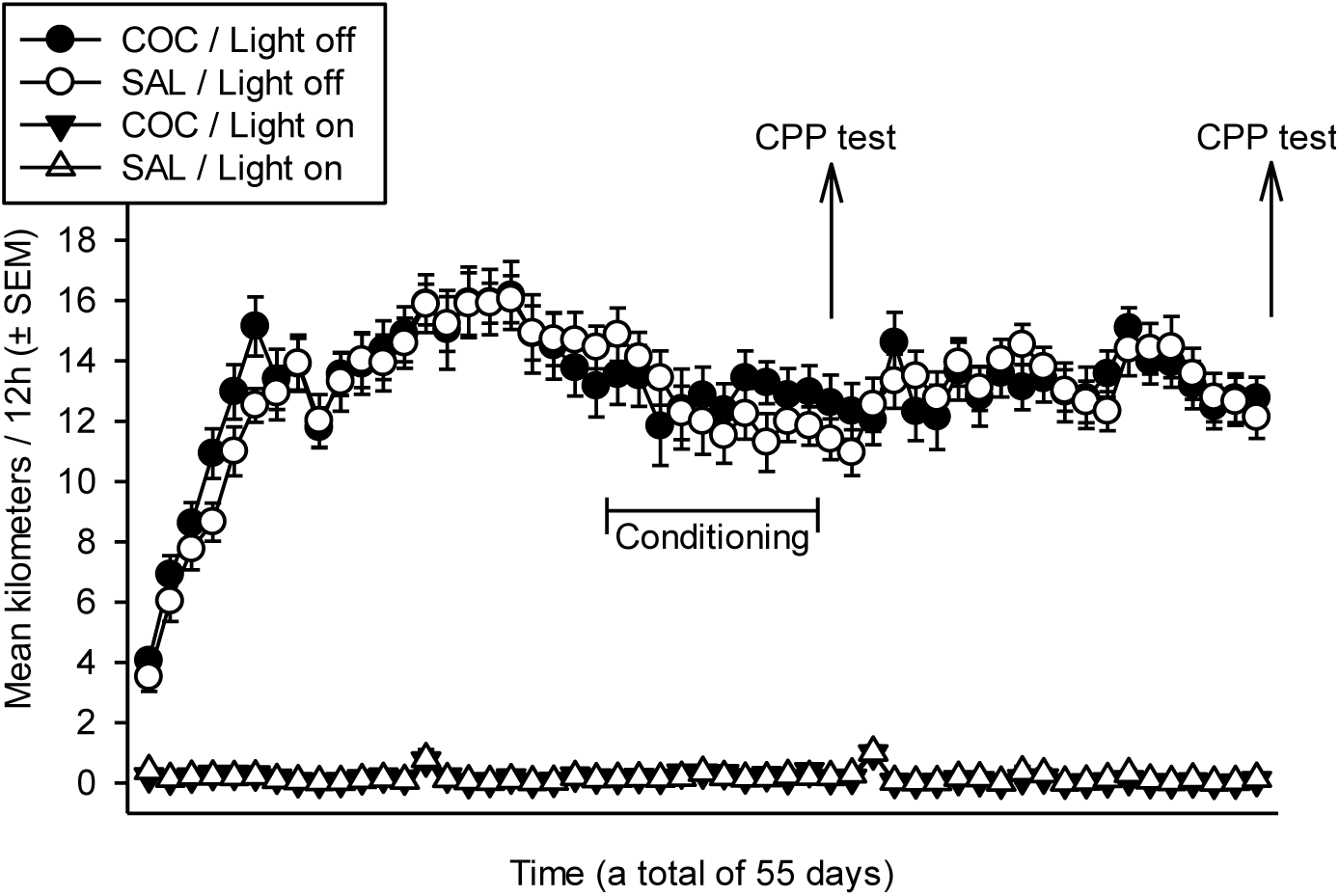
Running activity. Nocturnal (light off) and diurnal (light on) wheel-running exercise (distance traveled in kilometers).

Fig.3 presents CPP tested 1 day and 21 days after the last session of conditioning (panels A and B respectively). In the first preference test, “Wheel” and “No Wheel” cocaine groups exhibited similar levels of well-marked preference for the drug-paired compartment (A). Planned comparisons confirmed the much higher CPP scores of cocaine groups over those of their respective saline controls in both groups (“Wheel”: *d* = 1.38, *t*_(53)_ = 5.04, *p* < 0.001 one-tailed and “No Wheel”: *d* = 1.11, *t*_(53)_ = 4.04, *p* < 0.001 one-tailed). However, the effect underlying interaction between housing and the pharmacological treatment (resulted from the ANOVA) was clearly negligible (η²p = 0.009, F_(1,53)_ = 0.50, *p* = 0.48).

**Fig. 3.**
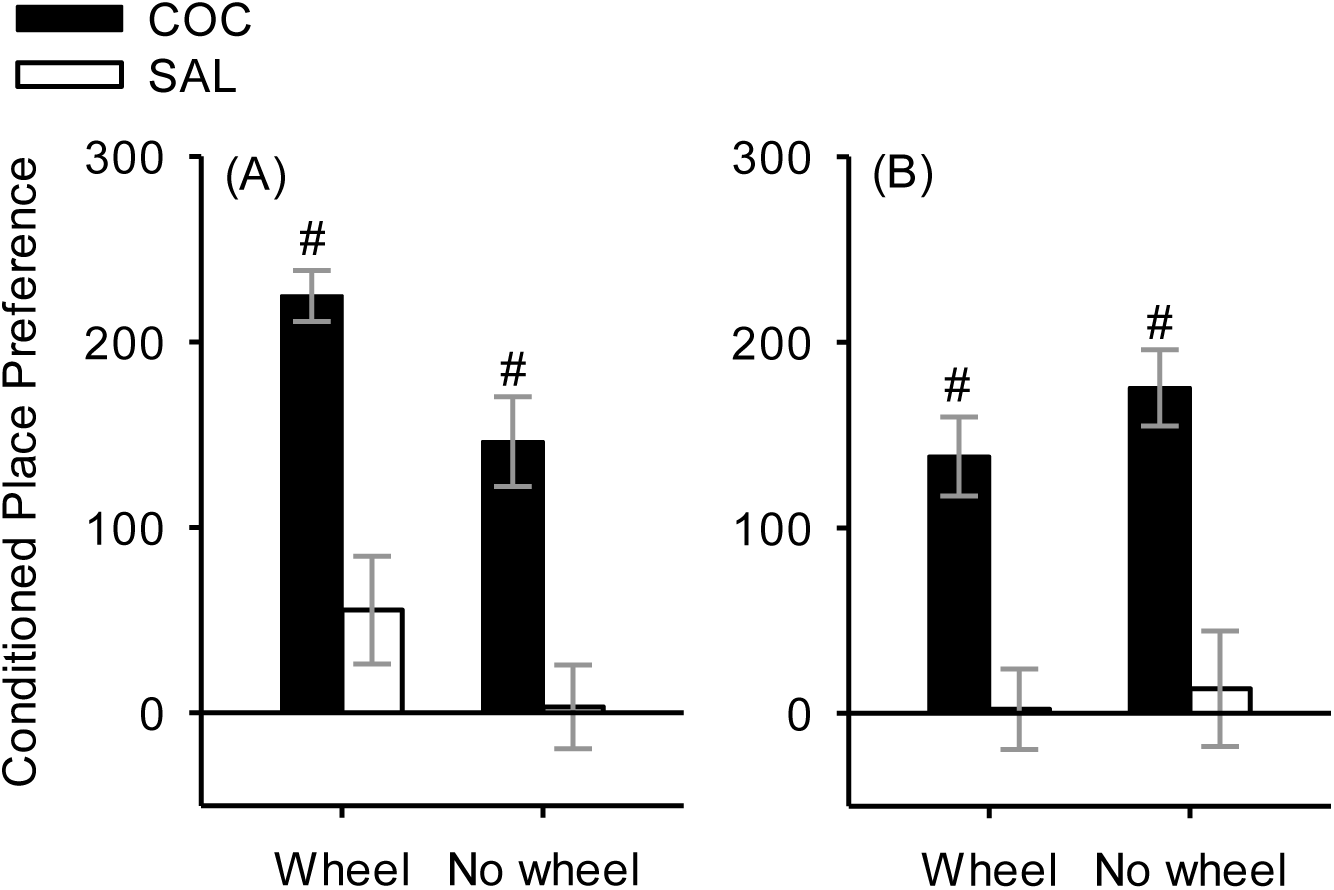
Conditioned place preference (CPP). CPP scores 1 day (A) or 21 days (B) after the last conditioning session. #: much higher than the corresponding saline group (minimal Cohen’s *d* at 1.09).

The clear-cut preference for the drug-paired compartment displayed by the two cocaine groups was still present 21 days after the last conditioning session (all mice being tested under saline). Planned comparisons revealed effect sizes comparable to those reported in the first CPP test (“Wheel”: *d* = 1.09, *t*_(53)_ = 3.95, *p* < 0.001 one-tailed and “No Wheel”: *d* = 1.15, *t*_(53)_ = 4.19, *p* < 0.001 one-tailed). However, the obtained data were far from being incompatible with the model prediction under the (null) hypothesis, that is an absence of interaction (η²p < 0.005, F_(1,53)_ = 0.028, *p* = 0.87).

These results reproduce those previously obtained in our lab [9] by showing that wheel-running exercise is ineffective at convincingly altering CPP to 10 mg/kg cocaine in male C57BL/6J mice (negligible effect found at ƞ²p ≤ .01 in both studies). Despite different procedural parameters such as the interval between housing assignment and testing (21 vs. 70 days) and apparatus (two vs. three compartments), mice receiving cocaine exhibited strongly-marked CPP scores in comparison to their saline controls. Here, these clear-cut effects were observed regardless of the post-conditioning interval (1 day or 21 days), highlighting the robustness of that phenomenon (cocaine reward). Note that comparable long-term expression of cocaine-induced CPP was previously observed in male C57BL6 tested in a drug-free state 28 days after place preference conditioning [11].

The lack of evidence for a sizeable difference in cocaine-induced CPP between exercised and sedentary individuals is in agreement with results reported with a low dose of 3 mg/kg morphine in rats housed with a wheel for 42 days before testing [8]. While wheel-running blocked stress-induced accentuation of CPP to morphine, such regimen of exercise did not impact CPP per se (in rats that were not submitted to the stress procedure consisting of inescapable shocks).

The results from the two other studies that have investigated the effects of wheel-running on CPP induced by cocaine contrast with ours by reporting accentuated CPP after voluntary exercise [6,7]. In Smith and colleagues [6], Long–Evans female rats were conditioned to 0 (control), 5 or 10 mg/kg cocaine after being housed with or without a running wheel from 21 to 63 days of age (6 weeks). Although the interaction between dose and housing condition was not significant, exercise resulted in an increase in time spent into the compartment previously paired with 10 mg/kg cocaine as compared to sedentary rats (detected from multiple pairwise comparisons). In the Mustroph and collaborators [7], following a 30-day period of housing with a running wheel from around 54 to 84 days of age, C57BL/6J mice receiving 10 mg/kg cocaine exhibited greater CPP scores than those displayed by the sedentary mice. However, in these studies, the pharmacological control was separately analyzed, and therefore the strength of the interaction between the drug treatment (and the dose) and housing condition was unclear. That said, their results are convergent in spite of critical procedural differences such as species, ages, and durations of wheel access. This exercise-induced increase in CPP somewhat contrasts with the protective effects of aerobic exercise on vulnerability to the rewarding properties of drugs of abuse [12].

Such an accentuation in conditioned reward to drugs after free exercise has been hypothetically ascribed to an enhancement of associative learning capabilities resulting from neuroplastic changes induced by wheel-running [7,13-15]. This possibility is further supported in the Mustroph and colleagues’ study [7] by the increase of the formation of CPP and its delayed extinction when wheel-running took place before acquisition, whereas extinction was facilitated when exercise was administered after acquisition. In that framework, one can speculate that the hypothetical inhibition of wheel-running of the rewarding component of CPP in our mice was concurrently reversed by the improving effect of such exercise on contextual learning (that underlies CPP), finally leading to neutralized effects. One can further speculate that attenuated or augmented CPP scores in animals housed with a running wheel would depend on the relative strength of pro-cognitive and attenuating effects of the rewarding properties of drugs; this interaction may be modulated by experimental parameters. These results, along with others, question the nature and the robustness of the effects that exercise may induce on cocaine CPP.

## Acknowledgements

Louis-Ferdinand LESPINE was under contract (doctoral fellow) with the Fund for Scientific Research – FNRS (Belgium) during the realization of the present work.

